# Computational modelling of functional maturation of primary motoneuron firing properties in developing zebrafish

**DOI:** 10.1101/2025.10.17.683092

**Authors:** Stephanie F. Gaudreau, Tuan V. Bui

## Abstract

Several ion currents of zebrafish primary motoneurons undergo changes in expression level during early development. Similarly, the firing properties of primary motoneurons and their involvement during locomotor activity change during early development as locomotor control of developing zebrafish matures. To test whether the experimentally observed changes in ion currents during development could underlie changes in firing properties and in participation during locomotor activity, we created models of primary motoneurons at developmental stages. Changes in the expression levels of a persistent outward potassium current, persistent inward potassium current, and several high-voltage activated calcium currents were modelled based on experimental observations. Simulations of our computational models replicated shifts in primary motoneuron firing properties and involvement during light-evoked swimming observed at 3 and 5 days post-fertilization. Our results suggest that developmental changes in specific ion currents of primary motoneurons could be sufficient to foster changes in firing properties of primary motoneurons that shape their activity level during maturation of motor control in developing zebrafish.

**Key points:**

- Developmental changes in persistent inward and outward currents could explain changes in firing properties of primary motoneurons in larval zebrafish
- Modelling suggests these currents are sufficient to explain differences in primary motoneuron firing during light-evoked swimming responses at two developmental stages
- Developmental changes in high-voltage activated calcium currents explain differences in appearance of persistent inward currents during voltage-clamp ramp
- Interaction between high-voltage activated calcium currents and calcium-dependent potassium currents could explain why blocking calcium currents increases primary motoneuron firing

## INTRODUCTION

The firing properties of neurons can change during development (McGuirt *et al*., 2021; Feng *et al*., 2022; Mermet-Joret *et al*., 2022; Mohammadalinejad *et al*., 2024). These changes in firing behaviour can arise from developmental changes in intrinsic properties and active conductances expressed by neurons (Carlin *et al*., 2008; Ehrlich *et al*., 2012; Watanabe *et al*., 2017; Sánchez-Aguilera *et al*., 2020; Sharples & Miles, 2021; Singh *et al*., 2025). Considering the nonlinear operation of ion currents, ascribing specific changes in firing properties to developmental changes of specific ion currents can be difficult.

Developing zebrafish exhibit relatively rapid motor maturation at early stages. In the first days of development, movements rapidly emerge at the same time that new neurons are born and integrated into new motor circuits that span spinal, supraspinal and neuromuscular regions of the body (Myers *et al*., 1986; Drapeau *et al*., 2002; Lewis & Eisen, 2003; Roussel *et al*., 2021). Zebrafish will transition from movements such as coiling that are mainly characterized by ballistic, large-amplitude, full-body contractions to more mature swimming forms characterized by high-frequency, more subtle tail bends (Saint-Amant & Drapeau, 1998). Primary motoneurons (pMNs), the earlier-born subset of motoneurons, which innervate superficial, fast fatigable muscle fibers, are prominent during the early developmental stages when coiling and escape movements predominate but give way to the later-born secondary motoneurons that innervate deeper, fatigue-resistant, slow muscle fibers (Liu & Westerfield, 1988). Secondary motoneurons predominate during the swimming movements that are more prominent at later stages of development and into adulthood (McLean *et al*., 2007; Ampatzis *et al*., 2013). As the motor repertoire of zebrafish expands and pMNs become relatively less involved in overall motor activity, their firing properties also change (Gaudreau & Bui, 2024). pMNs will show greater levels of sustained firing and less spike frequency adaptation in response to depolarizing current steps.

Several ion currents expressed within spinal neurons have been shown to shape their firing properties and motor output (Buss *et al*., 2003; Tong & McDearmid, 2012; Song *et al*., 2020; Gaudreau & Bui, 2025*a*). We previously showed that several ion currents in pMNs change expression levels between 2-5 days post-fertilization (Gaudreau & Bui, 2024, Gaudreau & Bui, 2025*b*, Gaudreau & Bui, 2025*c*). For example, the persistent outward current known as the M-current (*I*_M_) increases between 2 and 3 dpf. This is a short-lived increase as *I*_M_decreases already at 4 and 5 dpf (Gaudreau & Bui, 2024). In contrast, the persistent inward current known as the persistent sodium current (*I*_Na,P_) decreases between 2 and 5 dpf (Gaudreau & Bui, 2025*b*). Some other currents, such as the high-voltage activated (HVA) L-type Ca^2+^ current (*I*_Ca,L_) showed no developmental changes, whereas another HVA, the P/Q-type Ca^2+^ current, emerged after 3 dpf (Gaudreau & Bui, 2025*c*). Our previous studies suggested that some of these developmental dynamics could explain the diminishing involvement of pMNs in the overall locomotor repertoire of developing zebrafish. For example, pMNs were often recruited during swimming evoked by sudden light-illumination at 3 but not at 5 dpf (Gaudreau & Bui, 2025*b*).

In this study, we create the first models of developing pMNs with a full suite of active conductances to test whether developmental dynamics of ion currents could underlie the changes in firing properties and behaviour that were observed experimentally. Our models replicate different firing responses to current steps observed between 3 and 5 dpf, changes in pMN involvement in light-evoked swimming at 3 versus 5 dpf, and the emergence of two persistent inward current peaks during voltage-clamp ramps after 3 dpf. Furthermore, we test whether interactions between HVA calcium currents and calcium-dependent potassium currents could underlie the paradoxical effects of blocking HVA currents, which lead to increased firing in pMNs (Gaudreau & Bui, 2025*c*).

## METHODS

We built a simplified model of a primary motoneuron consisting of a single cylindrical compartment with a length and diameter of 25 µm, at all ages modelled. The value of the specific resistivity of the cytoplasm (*R*_*i*_) selected for this study is 35.4 Ω/cm. The specific membrane capacitance (C_m_) was set to 1 μF/cm^2^ (Hille, 2001), and therefore, the total cell capacitance was 19.6 pF. We chose to keep cell dimensions constant throughout the ages modelled and therefore, to obtain input resistance values comparable to experimental data (Gaudreau & Bui, 2024), we set passive membrane resistance to 9,817 Ω/cm^2^ for the 3 dpf model, and 8,925 Ω/cm^2^ for the 5 dpf model.

The following ion currents were incorporated into the soma of primary motoneurons.

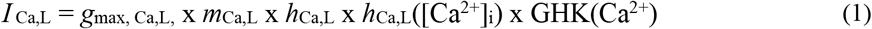

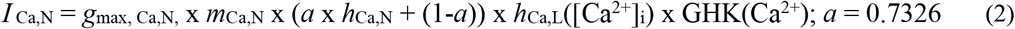

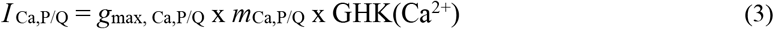

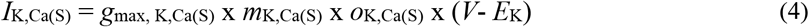

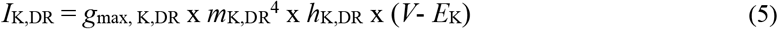

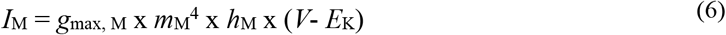

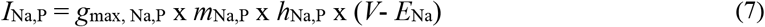

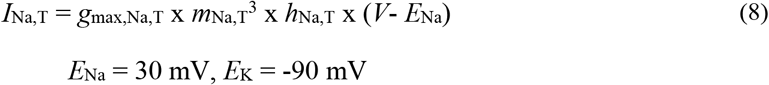

Where for current *i, g*_max,*i*_represents the peak conductance; and the variables *x*_i_are either the activation and inactivation variables, respectively, of the current *i*. GHK represents the Goldman-Hodgkin-Katz equation for ion flux (Hille, 2001), and *E*_Na_, *E*_K_, and *E*_Ca_represent the reversal potential for sodium and potassium currents, respectively.

For the strictly voltage-dependent active currents, the activation or inactivation variables are described by the following differential equation:

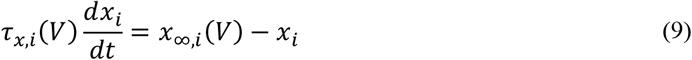

where *x* is the activation or inactivation variable of the current *i* (Table 1).

**TABLE 1.** Steady-state activation and inactivation variables and time constants for voltage-dependent currents.

| Ionic currents | Activation and inactivation variables | Time constants (in ms): |
| --- | --- | --- |
| $I_{Ca,L}$ | $m_{\infty, Ca,L} = [1 + \exp(-(V - (8.5))/4.3)]^{-1}$<br>$h_{\infty, Ca,L} = [1 + \exp((V - (-42.5))/(7.5))]^{-1}$<br>$h_{Ca,L}([Ca^{2+}]_i) = [1 + ([Ca^{2+}]/0.001)^4]$ | $\tau_{m, Ca,L} = 2.1 + [3.9 (\exp(-2(V - (-10))/16.0))^2]$<br>$\tau_{h, Ca,L} = 825.8 + [638.1 (\exp(-2(V - 0)/39.7))^2]$ |
| $I_{Ca,N}$ | $m_{\infty, Ca,N} = [1 + \exp(-(V - (-6.5))/6.5)]^{-1}$<br>$h_{\infty, Ca,N} = [1 + \exp((V - (-70))/(12.5))]^{-1}$ | $\tau_{m, Ca,N} = 0.8 + [5.4 (\exp(-2(V - (-20))/15.0))^2]$<br>$\tau_{h, Ca,N} = \text{Look-up table: } [(-30, 60), (-20, 50), (-10, 44.9), (0, 21.1), (10, 22.3), (20, 26.9)]$ |
| $I_{Ca,P/Q}$ | $m_{\infty, Ca,P/Q} = [1 + \exp(-(V - (-10.1))/3.1)]^{-1}$<br>$m_{\infty, Ca,P/Q} = [1 + \exp(-(V - (-45.1))/3.1)]^{-1}$<br>(for Fig. 3G) | $\tau_{m, Ca,P/Q} = 0.35 + [5.5 (\exp(-2(V - (-14.7))/18.1))^2]$<br>$\tau_{m, Ca,P/Q} = 0.35 + [5.5 (\exp(-2(V - (-49.7))/18.1))^2]$<br>(for Fig. 3G) |
| $I_{K,DR}$ | $m_{\infty, K,DR} = [1 + \exp(-(V - (-35))/7)]^{-0.25}$<br>$h_{\infty, K,DR} = 0.9 + 0.1/(1 + \exp((V - (-71))/150))$ | $\tau_{m, K,DR} = (120/(6 \exp(V - (-80))/24) + 70 \exp(-(V - (-59))/7)) + 1.3$<br>$\tau_{h, K,DR} = (1000/(\exp(V - (-60))/20) + \exp(-(V - (-60))/8)) + 50$ |
| $I_M$ | $m_{\infty, M} = [1 + \exp(-(V - (-19))/10)]^{-0.25}$<br>$h_{\infty, M} = 0.9 + 0.1/(1 + \exp((V - (-61))/150))$ | $\tau_{m, M} = (320/(4 \times \exp(V - (10))/70) + 10 \exp(-(V - (-53))/6)) + 10$<br>$\tau_{h, M} = (1000/(\exp(V - (-50))/20) + \exp(-(V - (-60))/8)) + 50$ |
| $I_{K,Ca(S)}$ | $n_{\infty, K,Ca(S)} = [1 + \exp((V - (E_K - V_{1/2})/sf)]^{-1}$<br>$o_{\infty, K,Ca(S)} = [Ca^{2+}]_i^{5.6}/(0.00042^{5.6} + [Ca^{2+}]_i^{5.6})$<br>$V_{1/2} = \text{Look-up table: } [(0.0002, 24), (0.0003, 35.3), (0.0005, 60.5), (0.001, 59.1)]$<br>$sf = \text{Look-up table: } [(0.0002, 128), (0.0003, 38.8), (0.0005, 48.8), (0.001, 44.8)]$ | |
| $I_{Na,P}$ | $m_{\infty, Na,P} = [1 + \exp(-(V - (-48))/3.1)]^{-1}$<br>$h_{\infty, Na,P} = [1 + \exp(-(V - (-40))/(-8))]^{-1}$ | $\tau_{m, Na,P} = 1$<br>$\tau_{h, Na,P} = 2000/\cosh((V - (-40))/16)$ |
| $I_{Na,T}$ | $m_{\infty, Na,T} = [1 + \exp(-(V - (-29))/8)]^{-1}$<br>$h_{\infty, Na,T} = [1 + \exp((V - (-53))/7)]^{-1}$ | $\tau_{m, Na,T} = (70/(5 \exp(V - (-100))/16) + 200 \exp(-(V - (-40))/10)) + 0.01$<br>$\tau_{h, Na,T} = (110/(3 \exp(V - (-100))/20) + 10 \exp(-(V - (-80))/3)) + 0.1$ |

Where, for the current i, *x*_∞,i_ (*V*) is the voltage-dependent steady-state activation or inactivation, respectively; and τ_*x*, i_(*V*) is the voltage-dependent time constant of the variable *x*. Modelling of currents in our models are based on prior models (Rybak *et al*., 2006; Bui & Brownstone, 2015; Watanabe *et al*., 2017; Mandge & Manchanda, 2018).

Calcium diffusion was modeled as described in previously (Booth *et al*., 1997). Internal calcium concentration is calculated from the following differential equation:

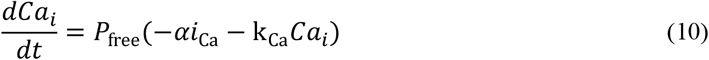

where *P*_free_is the percentage of free calcium to bound calcium and was set to 0.001; α is the total calcium current and has a value of µm/C, k_Ca_is the calcium removal rate and was assigned a value of 2/ms, and *i*_Ca_is the total calcium current. The steady-state internal calcium concentration was set at 1 x 10^-5^ mM.

### Neuron simulator

The models were created and simulations were run using NEURON v8.2.6+. The code to create the simulation environment, as well as the starting point for our primary motoneuron model, came from code to simulate Mauthner cells (Watanabe *et al*., 2017).

## RESULTS

### Changes in I_M_and I_Na,P_are sufficient to replicate changes in sustained spiking and spike frequency adaptation in pMNs during development

We have previously shown that the ability of pMNs to repetitively fire increases during early development. At 3 dpf, repetitive firing is limited, with an average of 8.33 ± 6.85 spikes generated in response to a current injection equal to 2x rheobase while 5 dpf pMNs have a greater capacity for repetitive firing with an average of 24.38 ± 12.16 spikes (Gaudreau & Bui, 2025*b*). This firing maturation coincides with developmental changes in the amplitude of *I*_M_, with its magnitude peaking at 3 dpf (42.88 ± 18.44 pA) and subsequently decreasing by 5 dpf (28.63 ± 11.56 pA) (Gaudreau & Bui, 2024). At the same time, we find that *I*_Na,P_decreases gradually in amplitude from 2 to 5 dpf (Gaudreau & Bui, 2025*b*). We aimed to replicate these developmental changes of *I*_M_and *I*_Na,P_in our simulated pMNs at 3 and 5 dpf to test whether the respective changes in the magnitudes of these opposing currents alone could explain maturation of repetitive firing in pMNs.

With parameter values for pMNs at each age described in Table 2, our modelled pMNs at both ages have membrane capacitances equaling 19.6 pF, which is similar to the experimental values previously reported (19.35 ± 3.05 pF at 3 dpf and 19.89 ± 3.268 pF at 5 dpf) (Gaudreau & Bui, 2024). Input resistance, calculated by membrane potential changes to −10 pA injections to pMN models held at approximately −65 mV, was measured at 350 MΩ and 330 MΩ from our simulated 3 and 5 dpf pMNs, respectively, which is also similar to experimental values previously reported (340.2 ± 103.6 MΩ at 3 dpf and 292.5 ± 192.1 MΩ at 5 dpf) (Gaudreau & Bui, 2024). With conductance values for *I*_Na,T_, *I*_K,DR_, *I*_M_and *I*_Na,P_as listed in Table 2, we find that 3 dpf simulated pMNs generate 11 spikes while 5 dpf simulated pMNs generate 39 spikes in response to a 2x rheobase current injection (**Fig. 1*A,D***). Rheobase values in simulated pMNs were similar to those observed experimentally with values of 70 pA and 100 pA at 3 and 5 dpf respectively (3 dpf: 75.00 ± 26.13 pA and 5 dpf: 94.77 ± 31.49 pA) (Gaudreau & Bui, 2024).

**TABLE 2.**
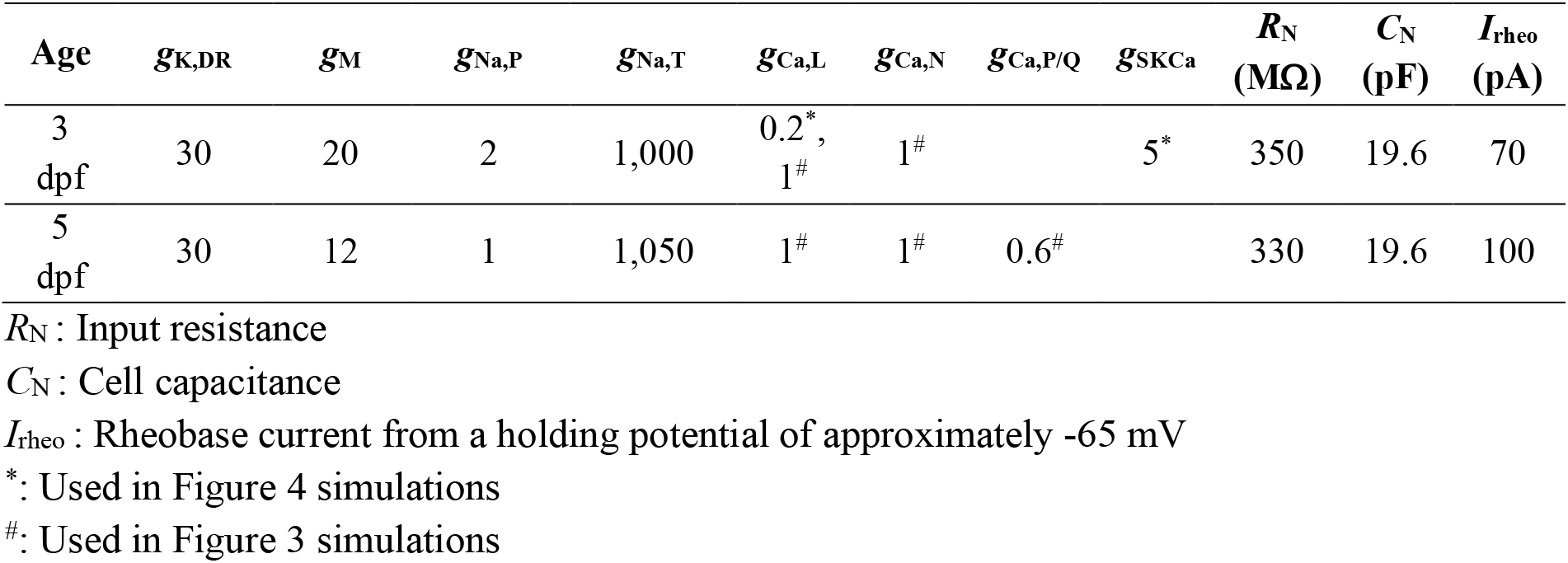
Maximum conductance values (nS/soma area) and intrinsic properties of pMN models at different ages.

| Age | $g_{K,DR}$ | $g_M$ | $g_{Na,P}$ | $g_{Na,T}$ | $g_{Ca,L}$ | $g_{Ca,N}$ | $g_{Ca,P/Q}$ | $g_{SKCa}$ | $R_N$<br>(MΩ) | $C_N$<br>(pF) | $I_{rheo}$<br>(pA) |
| --- | --- | --- | --- | --- | --- | --- | --- | --- | --- | --- | --- |
| 3<br>dpf | 30 | 20 | 2 | 1,000 | 0.2<sup>*</sup>,<br>1<sup>#</sup> | 1<sup>#</sup> |  | 5<sup>*</sup> | 350 | 19.6 | 70 |
| 5<br>dpf | 30 | 12 | 1 | 1,050 | 1<sup>#</sup> | 1<sup>#</sup> | 0.6<sup>#</sup> |  | 330 | 19.6 | 100 |

$R_N$ : Input resistance
$C_N$ : Cell capacitance
$I_{rheo}$ : Rheobase current from a holding potential of approximately -65 mV
\*: Used in Figure 4 simulations

**Figure 1.**
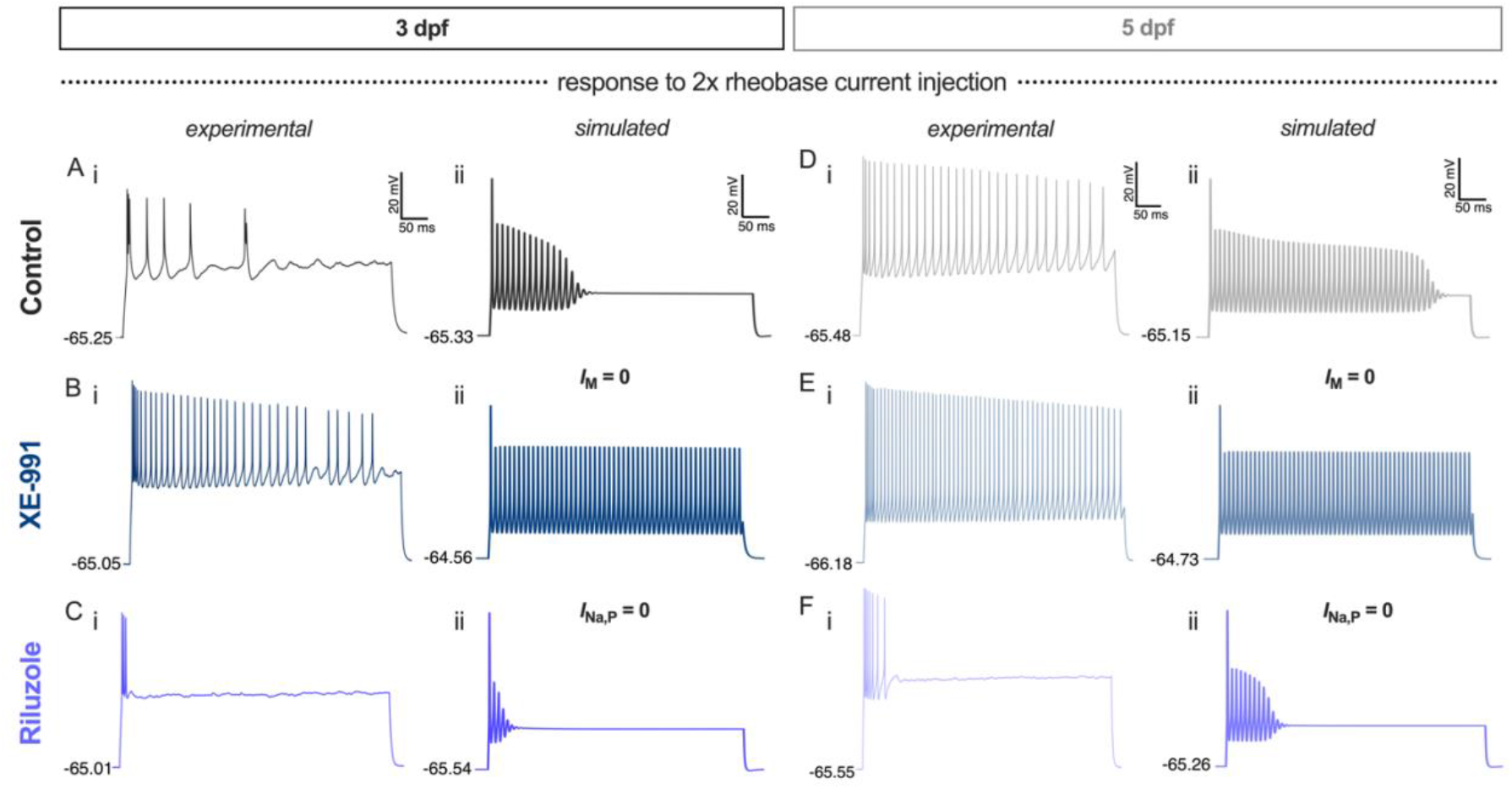
Simulated changes in *I*_M_and *I*_NaP_magnitudes replicate experimentally observed changes to pMN firing during development. Representative voltage responses obtained experimentally as previously described (Gaudreau & Bui, 2024). ***A***, Representative voltage response to a current injection equal to 2x rheobase from (*i*) a control pMN obtained experimentally and (*ii*) simulated pMN at 3 dpf. ***B***, Representative voltage response to a current injection equal to 2x rheobase from (*i*) a pMN exposed to 10 μM XE-991 and (*ii*) simulated pMN with *I*_M_= 0 at 3 dpf. ***C***, Representative voltage response to a current injection equal to 2x rheobase from (*i*) a pMN exposed to 5 μM riluzole and (*ii*) simulated pMN with *I*_Na,P_= 0 at 3 dpf. ***D***, Representative voltage response to a current injection equal to 2x rheobase from (*i*) a control pMN obtained experimentally and (*ii*) simulated pMN at 5 dpf. ***E***, Representative voltage response to a current injection equal to 2x rheobase from (*i*) a pMN exposed to 10 μM XE-991 and (*ii*) simulated pMN with *I*_M_= 0 at 5 dpf. ***F***, Representative voltage response to a current injection equal to 2x rheobase from (*i*) a pMN exposed to 5 μM riluzole and (*ii*) simulated pMN with *I*_Na,P_= 0 at 5 dpf. ***A****-****C***, a holding current of 55 pA was applied to the models. ***D****-****F***, a holding current of 60 pA was applied to the models.

When *I*_M_was inhibited experimentally by bath application of Kv7.2/7.3 *I*_M_-mediating channel blocker XE-991, repetitive firing was increased most prominently at 3 dpf, when the magnitude of *I*_M_is at its largest (Gaudreau & Bui, 2024). When we set *I*_M_= 0 in our simulated pMNs to replicate *I*_M_inhibition by XE-991, we find that the number of spikes generated in response to a 2x rheobase current injection increases to 53 and 58 at 3 and 5 dpf, respectively (**Fig. 1*B,E***). Setting *I*_Na,P_= 0 to replicate *I*_Na,P_inhibition by riluzole diminished the number of spikes generated more prominently at 3 dpf than at 5 dpf when compared to control simulations, as observed experimentally when pMNs were exposed to riluzole (**Fig. 1*C,F***) (Gaudreau & Bui, 2025*b*). Overall, our simulated pMN data demonstrates that the experimentally observed developmental changes in *I*_M_and *I*_NaP_may be sufficient to underlie the maturation of repetitive firing during early larval zebrafish development.

### Different pMN firing behaviours during light-evoked swimming in larval zebrafish could be due to differences in I_M_and I_NaP_

We hypothesized that these developmental changes in the magnitudes of *I*_M_and *I*_NaP_were responsible for observed differences in pMN recruitment during light-evoked escape responses at 3 versus 5 dpf. At 3 dpf, pMNs were readily recruited (**Fig. 2*A***) while pMNs at 5 dpf rarely were (**Fig. 2*E***) (Gaudreau & Bui, 2025*b*). We sought to simulate the response of pMNs at 3 and 5 dpf to a rhythmic semi-sinusoidal wave to mimic rhythmic locomotor drive received by pMNs during light-evoked swim responses. The current amplitude of the semi-sinusoidal wave is the same for 3 and 5 dpf pMNs as experimentally measured synaptic drive to 3 and 5 dpf pMNs was similar (Gaudreau & Bui, 2025*b*). We find that 3 dpf pMNs fire in response to the semi-sinusoidal wave, while 5 dpf pMNs do not in our simulations (**Fig. 2*C,G***). When *I*_M_= 0 to mimic inhibition of *I*_M_by XE-991, 3 dpf pMNs fire more spikes in response to the same semi-sinusoidal drive while 5 dpf pMNs remain unrecruited (**Fig. 2*D,H***), as seen in previous experimental recordings (**Fig. 2*B,F***) (Gaudreau & Bui, 2025*b*). These results suggest that the larger magnitude of *I*_Na,P_helps to recruit pMNs during light-evoked swim responses at 3 dpf and the lower magnitude of *I*_NaP_at 5 dpf limits the ability of pMNs to be recruited while complimentary magnitudes of *I*_M_help to ensure control over this recruitment.

**Figure 2.**
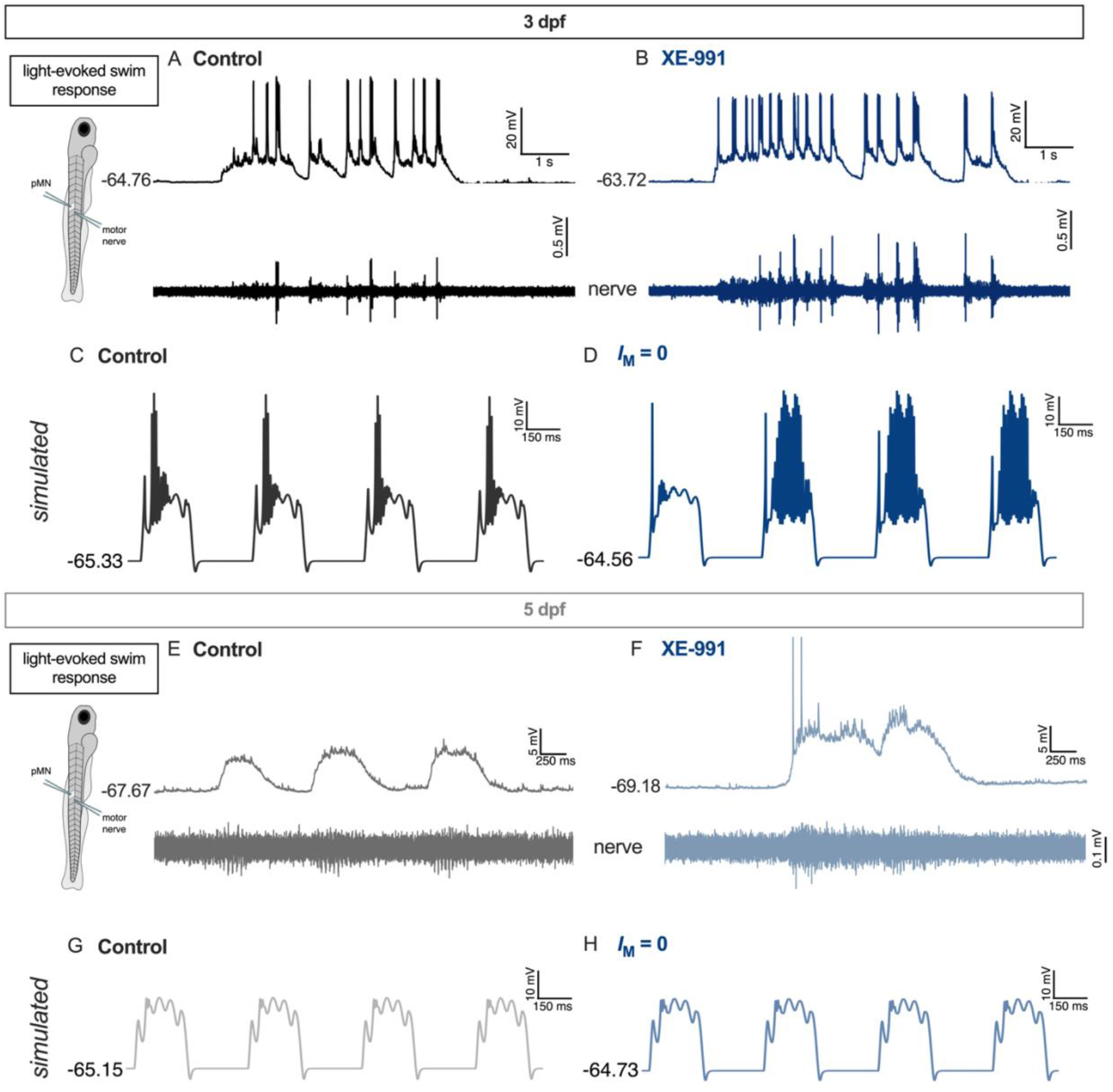
Simulated differences in *I*_M_and *I*_NaP_during development replicate pMN recruitment differences during light-evoked swim response at 3 and 5 dpf. ***A***, Representative voltage traces from a pMN (*top*) and motor nerve (*bottom*) during a light-evoked swim response at 3 dpf. ***B***, Representative voltage traces from a pMN (*top*) and motor nerve (*bottom*) exposed to 10 μM XE-991 during a light-evoked swim response at 3 dpf. ***C***, Voltage response from simulated 3 dpf pMN in response to a simulated rhythmic synaptic drive mimicking the drive evoked during light-evoked swim responses. ***D***, Voltage response from simulated 3 dpf pMN with *I*_M_= 0 in response to a simulated rhythmic synaptic drive mimicking the drive evoked during light-evoked swim responses. ***E***, Representative voltage traces from a pMN (*top*) and motor nerve (*bottom*) during a light-evoked swim response at 5 dpf. ***F***, Representative voltage traces from a pMN (*top*) and motor nerve (*bottom*) exposed to 10 μM XE-991 during a light-evoked swim response at 5 dpf. ***G***, Voltage response from simulated 5 dpf pMN in response to a simulated rhythmic synaptic drive mimicking the drive evoked during light-evoked swim responses. ***H***, Voltage response from simulated 3 dpf pMN with *I*_M_= 0 in response to a simulated rhythmic synaptic drive mimicking the drive evoked during light-evoked swim responses. ***C, D, G***, and ***H***, a periodic 180 pA, 2 Hz semi-sinusoidal wave, summed with 40 pA, 20 Hz sinusoidal waves on each crest were simulated.

### Contributions of HVA calcium currents to separate PICs in pMNs

We have previously described the contributions of high-voltage activated (HVA) calcium currents, including *I*_Ca,N_, *I*_Ca,L,_and *I*_Ca,PQ_, to persistent inward currents (PIC) measured in pMNs from 2 to 5 dpf (Gaudreau & Bui, 2025*c*). Interestingly, we revealed that current responses to a 20 mV/s voltage ramp starting at −90 mV displayed a second inward current (PIC 2) at more depolarized membrane potentials in pMNs only after 3 dpf (**Fig. 3*A,B***). The amplitude of the second PIC was reduced in pMNs having been exposed to ω-agatoxin, a potent P/Q-type calcium channel inhibitor (Gaudreau & Bui, 2025*c*). We wondered whether *I*_Ca,PQ_appeared as a second PIC due to depolarized voltage-dependency compared to *I*_Na,P,_*I*_Ca,L_,and *I*_Ca,N_, or whether this could be due to its dendritic expression further away from the recording site rather than somatic expression. We included *I*_Ca,L_and *I*_Ca,N_, in our single compartment pMN models at 3 and 5 dpf with *I*_Ca,PQ_included only at 5 dpf (**Fig. 3*C,D***). By setting the voltage-dependence of *I*_Ca,PQ_to more depolarized membrane potentials compared to the other currents that make up the PIC (see Table 1), our 5 dpf pMN model replicated the emergence of a second PIC when a 20 mV/s voltage ramp was applied (**Fig. 3*C,D***).

**Figure 3.**
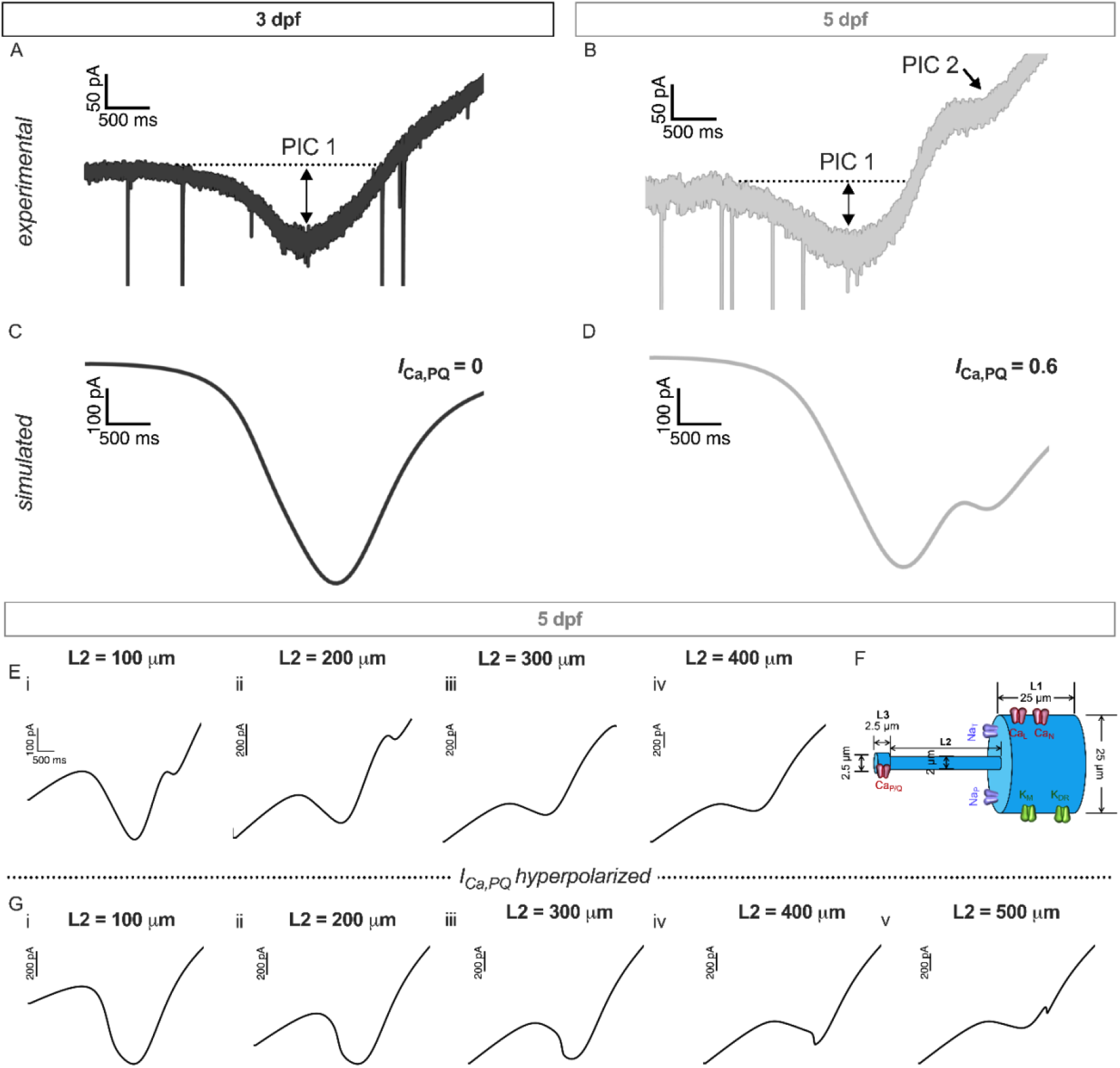
pMN simulations support emergence of somatic *I*_Ca,PQ_with depolarized voltage-dependency. ***A*-*B***, Representative current response containing a persistent inward current (PIC 1) to a 20 mV/s voltage ramp in a pMN at (***A***) 3 dpf and (***B***) 5 dpf. Note the appearance of a second persistent inward current (PIC 2) at 5 dpf. Traces shown previously (Gaudreau & Bui, 2025*c*). ***C***-***D***, Simulated current response to a 20 mV/s voltage ramp in a single compartment pMN model at (***C***) 3 dpf and (***D***) 5 dpf. ***E***, Simulated current responses to a 20 mV/s voltage ramp in a three-compartment pMN model when linking compartment L2 is modified to (*i*) 100 μm, (*ii*) 200 μm, (*iii*) 300 μm, and (*iv*) 400 μm. ***F***, Representation of three compartment pMN model. ***G***, Simulated current responses to a 20 mV/s voltage ramp in a three-compartment pMN model when L2 is modified to (*i*) 100 μm, (*ii*) 200 μm, (*iii*) 300 μm, (*iv*) 400 μm, and (*v*) 500 μm when *I*_Ca,PQ_voltage-dependency is hyperpolarized.

While PICs have been observed in many vertebrate spinal motoneurons and interneurons (Lee & Heckman, 1996; Li *et al*., 2004; Theiss *et al*., 2007; Dai & Jordan, 2010), seldom, if ever, do they manifest as two separate peaks. To test whether PIC 2 could arise from *I*_Ca,PQ_located in a neuritic location, we next simulated current responses to the same 20 mV/s voltage ramp in a three-compartment 5 dpf pMN model with *I*_Ca,PQ_expressed only in the furthest compartment, simulating dendritic *I*_Ca,PQ_(**Fig. 3*F***). We find that as the distance between the furthest compartment increases relative to the modelled soma, by varying the length of the linking compartment L2, the second PIC mediated by *I*_Ca,PQ_disappears (**Fig. 3*E***). We then asked whether a second PIC could arise from a modified *I*_Ca_,_PQ_that has a similar activation range as the other two HVA currents (see Table 1). We performed the same simulations in 5 dpf pMN models wherein the activation of *I*_Ca_,_PQ_is hyperpolarized to fall within the activation voltage range of the first PIC. As the length of L2 is increased, bringing *I*_Ca,PQ_further from the soma, we observe the emergence of a second PIC (**Fig. 3*G***). However, this second PIC had a very steep onset when compared to experimental recordings, suggesting an all-or-none activation, perhaps due to less effective voltage-clamping due to space-clamp limitations. Overall, these results demonstrate that while the second PIC could arise from dendritic *I*_Ca,PQ_, the shape of the experimentally observed second PIC suggests that it is more likely to be from a somatic or perisomatic *I*_Ca,PQ_.

### I_Ca,L_can dampen pMN firing through activation of I_K,Ca_

While *I*_*Ca,L*_has been associated with increased excitability in vertebrate motoneurons (Lee & Heckman, 1996; Booth *et al*., 1997; Bennett *et al*., 1998; Perrier & Hounsgaard, 2003), our recordings with the L-type Ca^2+^ channel blocker nifedipine revealed a paradoxical increase in pMN firing (**Fig. 4*A,B***) (Gaudreau & Bui, 2025*c*). To investigate whether this could be due to the activation of calcium-dependent potassium currents, we modified our 3-dpf model by the addition of the SK potassium channels, *I*_K,Ca(S)_(see Table 2 for conductance densities). We found that *I*_K,Ca(S)_could be recruited by Ca^2+^ entry through *I*_Ca,L_(**Fig. 4*E,G***). This was confirmed when we simulated the effects of nifedipine by setting *I*_Ca,L_to 0, which practically reduced *I*_K,Ca(S)_to 0 as well (**Fig. 4*F,H***). Consequently, the effects of eliminating *I*_K,Ca(S)_by virtue of blocking *I*_*Ca,L*_, increased pMN firing in our simulations (**Fig. 4*C,D***) as observed experimentally (**Fig. 4*A,B***).

**Figure 4.**
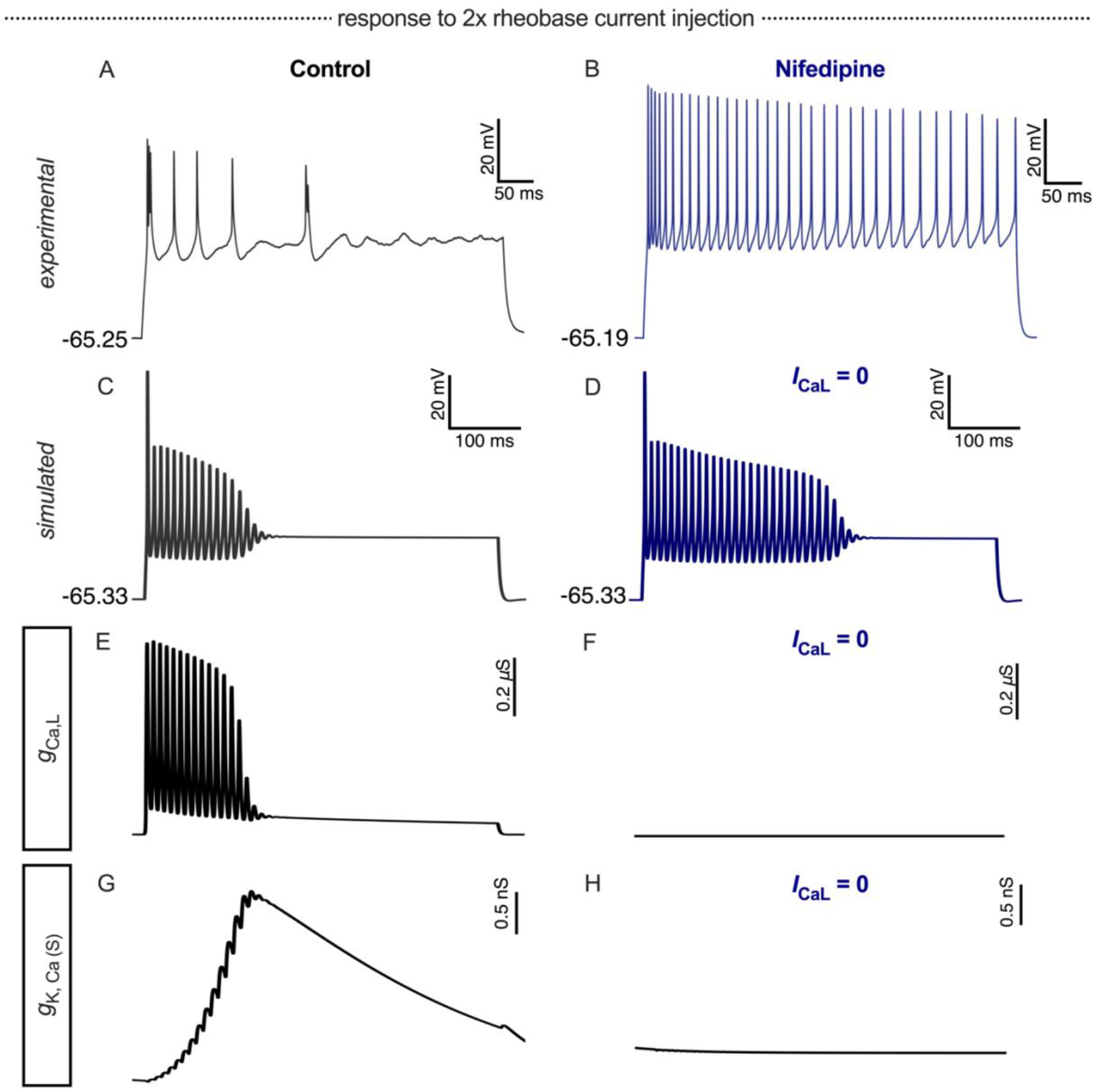
pMN simulations suggest that regulation of firing observed experimentally by *I*_Ca,L_at 3 dpf is linked to the activation of *I*_K,Ca(S)_. ***A***, Representative voltage response to a current injection equal to 2x rheobase from a control pMN obtained experimentally at 3 dpf. ***B***, Representative voltage response to a current injection equal to 2x rheobase from a pMN exposed to 20 μM nifedipine at 3 dpf. ***C***, Simulated 3 dpf pMN response to current injection equal to 2x rheobase. ***D***, Voltage response to 2x rheobase current injection from simulated 3 dpf pMN with *I*_Ca,L_= 0. ***E***, Conductance of *I*_Ca,L_during voltage response to 2x rheobase current injection in a simulated 3 dpf pMN. ***F***, Conductance of *I*_Ca,L_during voltage response to 2x rheobase current injection in a simulated 3 dpf pMN with *I*_Ca,L_= 0. ***G***, Conductance of *I*_K,Ca(S)_during voltage response to 2x rheobase current injection in a simulated 3 dpf pMN. ***F***, Conductance of *I*_K,Ca(S)_during voltage response to 2x rheobase current injection in a simulated 3 dpf pMN with *I*_Ca,L_= 0.

## DISCUSSION

The active conductances of developing zebrafish pMNs exhibit a mixture of developmental profiles (Gaudreau & Bui, 2024). Electrophysiological recordings suggest that while the persistent inward current, *I*_Na,P_, decreases between 2 and 5 dpf, the persistent outward current, *I*_M_, peaks transiently at 3 dpf (Gaudreau & Bui, 2025*b*). On the other hand, the HVA *I*_Ca,L_remains constant throughout the 2-5 dpf time window, whereas *I*_Ca,P/Q_emerges after 3 dpf (Gaudreau & Bui, 2025*c*). During this developmental timeline, the firing properties of pMNs also undergo transformation. pMNs increasingly exhibit reduced spike firing adaptation, increasing their ability to sustain firing during long current stimulation. However, their contributions to locomotor activity decrease as evidenced by reduced involvement during light-evoked swimming (Gaudreau & Bui, 2025*b*). Whether the developmental changes that have been observed in several ion currents could be directly linked to these changes in firing properties and behaviours in pMNs of developing zebrafish was tested using computational modelling.

### Decreased spike frequency adaptation following changes in persistent currents

Our models of 2-5 dpf pMNs incorporated progressively diminishing *I*_Na,P_and a transient *I*_M_peak at 3 dpf. We ensured that our models exhibited input resistance and rheobase values that were in line with experimental values (Gaudreau & Bui, 2024). With these electrophysiological conditions satisfied, our models exhibited spike numbers as well as levels of sustained spiking at various levels of current injections that were in line with experimental recordings (Gaudreau & Bui, 2025*b*). Also, the changes in these firing properties when either *I*_M_or *I*_Na,P_were blocked in our model were consistent with electrophysiological recordings of pMN during pharmacological modulation of these currents (Gaudreau & Bui, 2024, Gaudreau & Bui, 2025*c*, Gaudreau & Bui, 2025*b*). One additional factor that greatly influenced the ability to sustain firing was the current density of the fast transient sodium current. Our modelling work motivates future experimental investigation of this current in relation to firing behaviour in pMNs, and whether there are possible associations between changes in the currents that we have already investigated and changes in the fast, inactivating, sodium current.

### Developmental differences in pMN activity during light-evoked swimming could be shaped by changes in persistent currents

Our experimental data suggested that developmental changes in *I*_M_or *I*_Na,P_could change the participation of pMNs in motor behaviours of developing zebrafish such as light-evoked swimming (Gaudreau & Bui, 2025*b*). We simulated the locomotor drive received by pMNs during light-evoked swimming using a semi-sinusoidal current injection to our 3 and 5 dpf pMN models. Consistent with electrophysiological recordings of pMNs during light-evoked swimming (Gaudreau & Bui, 2025*b*), the 3 dpf pMN model showed a greater amount of activity during the locomotor drive as compared to the 5 dpf model in response to the same stimuli. The amplitude of the stimulus had to be significantly increased before the 5 dpf model became active. As both models showed similar resting membrane potentials, our simulations suggest that differences in *I*_M_and *I*_Na,P_expression levels could account for the different responses at 3 and 5 dpf. Considering that the outward *I*_M_current is decreased from 3 to 5 dpf while the fast, inactivating sodium current is increased in our model, our simulations suggest that the decrease in the persistent inward current, *I*_Na,P_, is the main culprit for decreasing the level of pMN activity during light-evoked swimming. Thus, while pMNs are more capable of sustaining firing at 5 dpf as compared to 3 dpf (Gaudreau & Bui, 2024), putatively due to decreases in *I*_M_and an increase in the fast, inactivating sodium current, a decrease in *I*_NaP_seems to be powerful enough to remove pMNs participation from light-evoked swimming at that age.

### The origins of the second PIC in pMNs

PICs in motoneurons have long been shown to manifest themselves as a downward deflection in the resulting current response during voltage-clamp ramps (Lee & Heckman, 1998; Perrier & Hounsgaard, 2003; Li *et al*., 2004; Brocard *et al*., 2013). These PICs appear around the threshold membrane potential for action potential firing and can be composed of a mix of *I*_Na,P_and *I*_CaL_. Our current recordings in 4-5 dpf pMNs in response to similar voltage-ramps have revealed a novel second downward deflection at more depolarized membrane potential values, which was attributed to *I*_Ca,P/Q_based on pharmacological blockade of this current (Gaudreau & Bui, 2025*c*). Our single-compartment model incorporated *I*_Ca,L_, *I*_Ca,N_and *I*_Ca,P/Q_. The difference in the activation profile of these currents, with the activation of *I*_Ca,L_and *I*_Ca,N_being more hyperpolarized than that of *I*_CaP/Q_, led to the appearance of two separate PICs.

*I*_Ca,P/Q_is often associated with the control of synaptic release, which suggests its presence at nerve terminals, including those of larval zebrafish (Wen *et al*., 2013). Our experimental data showing that blocking *I*_Ca,P/Q_with ω-agatoxin affects the firing of action potentials (Gaudreau & Bui, 2025*c*) suggests the additional presence of *I*_Ca,P/Q_at or near the cell body. We tested whether the second PIC could disappear if the location of *I*_Ca,P/Q_was moved away from the soma in a multi-compartmental model of a 5 dpf pMN. The second PIC did indeed disappear when moved far away enough from the soma. We then asked whether the second PIC could result from a *I*_Ca,P/Q_modified such that its activation properties were similar to *I*_Ca,L_and *I*_Ca,N_. The inclusion of this modified *I*_CaP/Q_at the soma resulted in a single PIC, due to the similar range of activation of all three HVA. When we moved this modified *I*_Ca,P/Q_further away from the cell body, we were eventually able to obtain a second PIC, but with an abrupt onset and steep downslope, which was never observed experimentally. This suggests an all-or-none activation of *I*_Ca,P/Q_, perhaps due to the larger distance away from our somatic voltage-clamp. These two sets of simulations and the disappearance of the second PIC when *I*_Ca,P/Q_is distanced from the cell body unless we modify its activation profile to be hyperpolarized, which leads to an abrupt all-or-none activation, provide supporting evidence that there may be a somatic *I*_Ca,P/Q_in pMNs in addition to its presence at the nerve terminal.

### Coupling of specific HVA calcium currents with calcium-dependent potassium currents

While HVA currents are often linked with increased neuronal excitability, such as the case of plateau potentials, pharmacological blockade of HVA currents in pMNs of developing zebrafish resulted in increased firing (Gaudreau & Bui, 2025*c*). This counterintuitive result could be due to the activation of calcium-dependent potassium currents. To test this possibility, we incorporated calcium-dependent potassium currents into our pMN models. At lower densities of HVA currents, the depolarization from the inward Ca^2+^ current was counteracted by the activation of calcium-dependent potassium currents. Removal of the inward Ca^2+^ current in our models replicated the increased pMN firing observed when HVA currents were blocked in pMNs. Note that the density of the *I*_Ca,L_for the simulations of the firing in response to step current injections was lower than for the simulations investigating the PICs observed during voltage-clamp ramps. The experimental recordings of PICs in pMNs were obtained using intracellular electrodes filled with cesium. Cesium improves the ability of voltage-clamp to control the membrane potential and may thus result in a greater activation of the L-type Ca^2+^ channels during voltage-clamp recordings than during current injections. Alternatively, cesium may reduce Ca^2+^ inactivation of the *I*_Ca,L_through a PKA-mediated pathway (Brette *et al*., 2003), and thus amplify its magnitude during our previous voltage-clamp recordings.

### Methodological considerations

By no means are our pMN models complete representations of their biological counterparts. While our simulations capture the essential features of the firing behaviour of pMNs and how they shift with developmental changes in ion currents modelled, the firing could be improved in terms of spike height and voltage-threshold. Improving these features could require refinements to the modelling of the currents or addition of new currents. Alternatively, a better morphological representation of the model could change firing properties. The single-model representation, while simple and computationally efficient, does not capture the dendritic and axonal complexities that are the hallmark of motoneurons (Rose *et al*., 1985; Bello-Rojas *et al*., 2019; Kishore *et al*., 2020). These neurite complexities are likely substrates for advanced synaptic integration (Rose & Cushing, 1999; Bui *et al*., 2008; Grande *et al*., 2010; Kishore *et al*., 2020) and could very well shape pMN firing properties (Mainen & Sejnowski, 1996). We also simplified intracellular calcium dynamics by omitting the contributions of mitochondrial and endoplasmic reticulum to calcium flow within cells (Moshkforoush *et al*., 2019). These dynamics may shape calcium-dependent activation and inactivation of the ion currents in our models, possibly coupling specific Ca^2+^ currents with specific Ca^2+^-dependent K^+^ currents (Viana *et al*., 1993; Tanabe *et al*., 1998; Goldberg & Wilson, 2005). We are also missing other ion currents expressed in pMNs, though we note the lack of LVA T-type Ca^2+^ currents or the hyperpolarization-activated inward current *I*_H_in pMNs as observed by the absence of any type of membrane depolarization activated by the application or termination of hyperpolarizing current injections in pMNs (Gaudreau & Bui, 2025*c*).

### Concluding remarks

We have built computational models of pMNs that capture experimentally observed changes in the expression profiles of several ionic conductances at two distinct stages during early zebrafish development. These models have allowed us to link specific changes in expression levels of ion currents to the maturation of firing and recruitment of pMNs during development. They also suggest likely sites of P/Q-type calcium current expression and potential interactions between HVA calcium currents and calcium-dependent potassium currents that may serve to limit firing in these neurons. Overall, this work provides mechanistic insight into how intricate developmental changes in ion conductances shape neuronal function.

## DATA AVAILABILITY

Models and data will be made available upon request.

## ACKNOWLEDGMENTS

This work was supported by the Natural Sciences and Engineering Research Council of Canada (NSERC): NSERC Discovery Grant, Grant Number: RGPIN-2022-03898 (to TVB); NSERC Canadian Graduate Scholarship M award, Award Number: NSERC 553401-2020 (to SFG); NSERC Postgraduate Scholarship D, Award Number: 569969-2022 (to SFG).

## AUTHOR CONTRIBUTIONS

SFG: Conceived and designed research, created models and performed simulations, analyzed data, interpreted results of simulations, prepared figures, drafted manuscript, edited and revised manuscript.

TVB: Conceived and designed research, created models and performed simulations, analyzed data, interpreted results of simulations, drafted manuscript, edited and revised manuscript, and approved final version of the manuscript.

